# Capillary stall quantification from optical coherence tomography angiogram maximum intensity projections

**DOI:** 10.1101/2021.10.01.461840

**Authors:** Signe K. Fruekilde, Eugenio G. Jiménez, Kim R. Drasbek, Christopher J. Bailey

## Abstract

Optical coherence tomography (OCT) is applicable to the study of cerebral microvasculature *in vivo*. Optimised acquisition schemes enable the generation of three-dimensional OCT angiograms, *i*.*e*., volumetric images of red blood cell flux in capillary networks, currently at a repetition rate of up to 1/10 seconds. This makes testable a new class of hypotheses that strive to bridge the gap between microscopic phenomena occurring at the spatial scale of neurons, and less invasive but crude techniques to measure macroscopic blood flow dynamics. Here we present a method for quantifying the occurrence of transient capillary stalls in OCT angiograms, *i*.*e*., events during which blood flow through a capillary branch is temporarily occluded. By making the assumption that information on such events is present predominantly in the imaging plane, we implemented a pipeline that automatically segments a network of interconnected capillaries from the maximum intensity projections (MIP) of a series of 3D angiograms. We then developed tools enabling rapid manual assessment of the binary flow status (open/stalled) of hundreds of capillary segments based on the intensity profile of each segment across time. The entire pipeline is optimized to run on a standard laptop computer, requiring no high-performance, low-availability resources, despite very large data volumes. To further reduce the threshold of adoption, and ultimately to support the development of reproducible research methods in the young field, we provide the documented code for scrutiny and re-use under a permissive open-source license.

## Introduction

The very small energy reservoirs of the brain place substantial requirements on the cerebral vasculature to constantly deliver the necessary oxygen and glucose to the cells of the brain. The mechanisms by which metabolic substrate delivery is matched to momentary energy needs due to neuronal signalling are collectively known as neurovascular coupling (reviewed in [1]). The majority of oxygen and nutrients diffuse from the blood to the neuronal tissue in the capillary bed, the vascular compartment of smallest-diameter vessels linking arterial and venous sides of the circulation. The capillaries make up most of the cerebral vasculature but are also its most fragile component. Microvascular dysfunction in the brain has been associated with neurodegenerative diseases and cognitive deficits (reviewed in [2-3]). Transient stalled flow in capillary segments [4] caused by neutrophil or platelet adhesions have been observed in various disease models [5-9], and may be an early component in the progression of cerebral microvascular dysfunction [2].

Studying the occurrence of stalled blood flow in the capillaries requires a longitudinal visualisation of the individual capillary segments. Although a two-photon microscope (TPM) can generate image sequences rapidly and with a very high signal-to-noise ratio (SNR), the number of capillaries captured in an imaging plane is limited [5-7]. Optical coherence tomography (OCT), while sacrificing some SNR and temporal resolution, allows for the visualisation of hundreds of capillary segments within a three-dimensional (3D) volume in consecutive ∼10-15 second frames [4, 8-9]. OCT images of brain vasculature are referred to as angiograms due to their visual similarity with radiographically acquired vessel maps. Despite the inherently low SNR of OCT, stalled blood flow leads to a robust reduction in the acquired signal and thus to high sensitivity. Imaging the occurrence of stall events thus amounts to a binary classification task on each capillary segment in a temporal sequence of angiograms.

Because capillary networks are inherently volumetric, the detection of stalls is a 3D problem that, at present, can only be performed manually. However, in our application of the technique, the optical properties of the imaging hardware limit the axial (in-depth) focus range of the 3D angiograms to 70–100 μm. We thus sought to develop software tools for a semi-automatic workflow in which capillary stalls are detected in two-dimensional (2D) maximum intensity projections (MIPs) of the acquired volumetric OCT angiograms. Here we provide a step-by-step walkthrough of the method, with illustrations of key analytic stages. We give examples of quantitative parameter estimates and their visualisation on capillary network maps. Limitations and potential future developments are discussed.

## Methods

Our analysis pipeline for OCT stall events is implemented in MATLAB (The MathWorks, Inc., USA), and draws heavily on open-source image processing routines made available by several authors. We have made every attempt to cite all relevant sources below. Our work is made available under the MIT License at https://github.com/cjayb/Fruekilde_OCT_stall_quant_paper.git, though we emphasise that elements of the code may be subject to more strict copyright clauses. The data used to demonstrate the pipeline is available for download on our Open Science Framework (OSF) project page: https://osf.io/me6hf/.

### Animal surgery and training

The data used to demonstrate the method were acquired in the piloting stage of an on-going study (in preparation), in which OCT data are acquired using the protocol described in [4, 8]. Seven C57BL/6J mice (N=7) had a chronic cranial window implanted and were trained to tolerate head fixation and remain calm during a single awake scanning session [10]. Sixty (60) consecutive 3D image frames were acquired in two non-overlapping regions-of-interest (ROI). The ROIs were acquired consecutively for a total imaging time of approximately 2 × 15 = 30 min. The data from one animal (WT02) were discarded due to excessive motion artefacts. Figures 1–4 were created using data from ROI1 of animal WT07.

**Figure 1.**
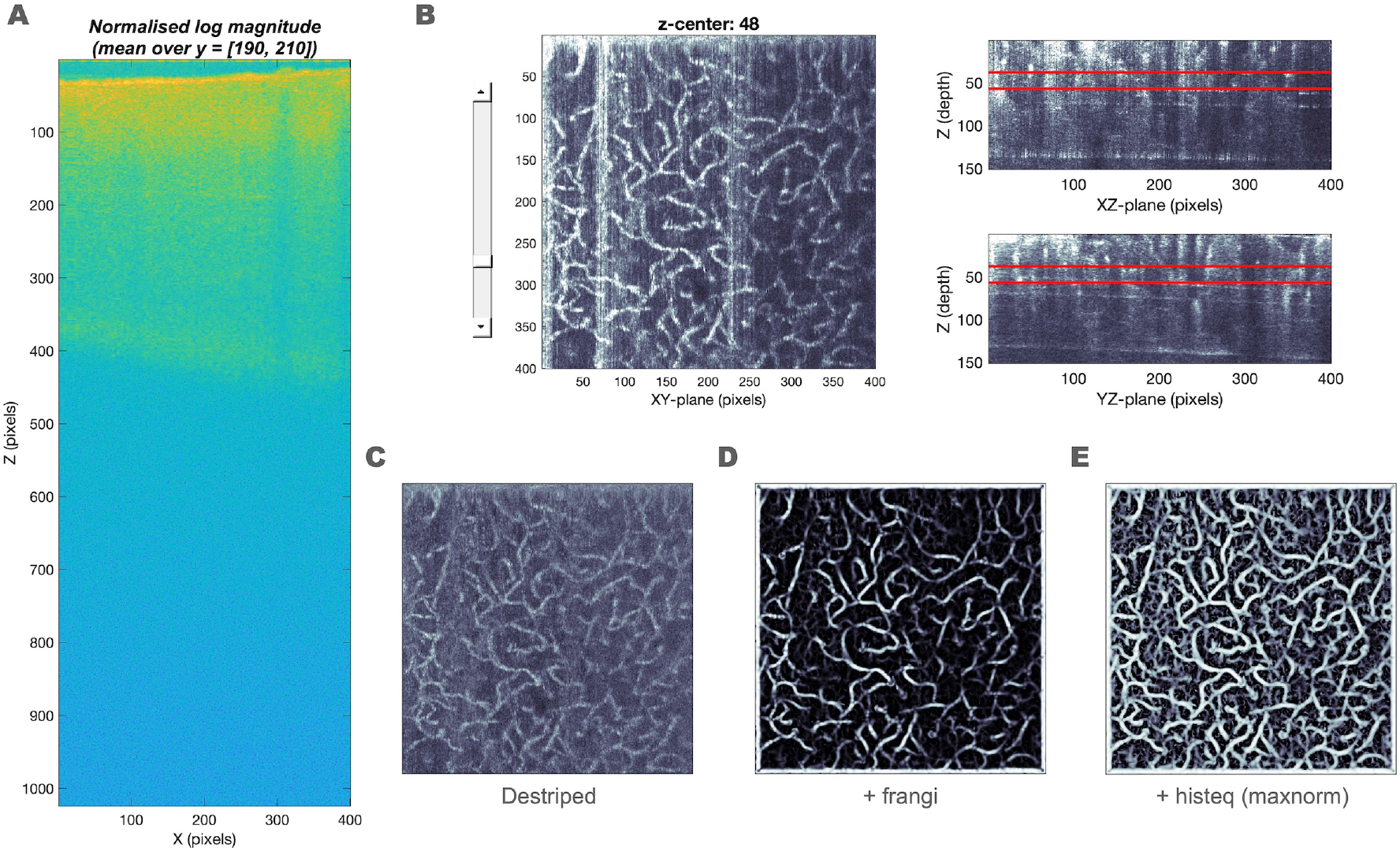
Reduction of Z dimension, image cleaning, and vessel extraction. **A)** The average of 20 Y-slices from the centre of the OCT collection (XZ plane). The focus plane is represented as the band of bright colour to depth of ca. 120 pixels. **B)** The GUI that enables the user to define the best focus plane. The image to the left shows the XY-plane MIP with a scroll bar to move in the Z-plane. The red lines mark the defined Z-stack (here 20 voxels deep) on the two images on the right. **C)** MIP from B after wavelet-based vertical de-striping. **D)** MIP from C after vessel-emphasising filter. **E)** MIP from D after contrast enhancement.

**Figure 2.**
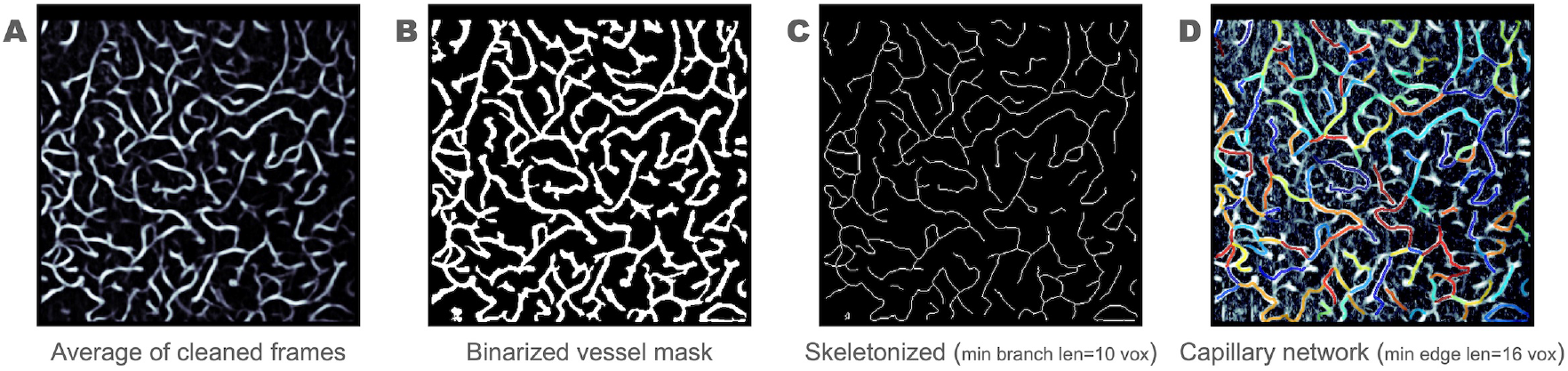
Identification of individual capillary segments. **A)** 2D images of the capillary network as an average of all 60 frames. **B)** Binary mask of the capillary network from A, with segments smaller than 100 pixels omitted. **C)** Skeletonized mask of the capillary network from B, with branches shorter than *25* μm omitted. **D)** Capillary segments (N=149) illustrated in different colours and overlaid on the averaged angiogram recomputed after correcting for in-plane motion.

**Figure 3.**
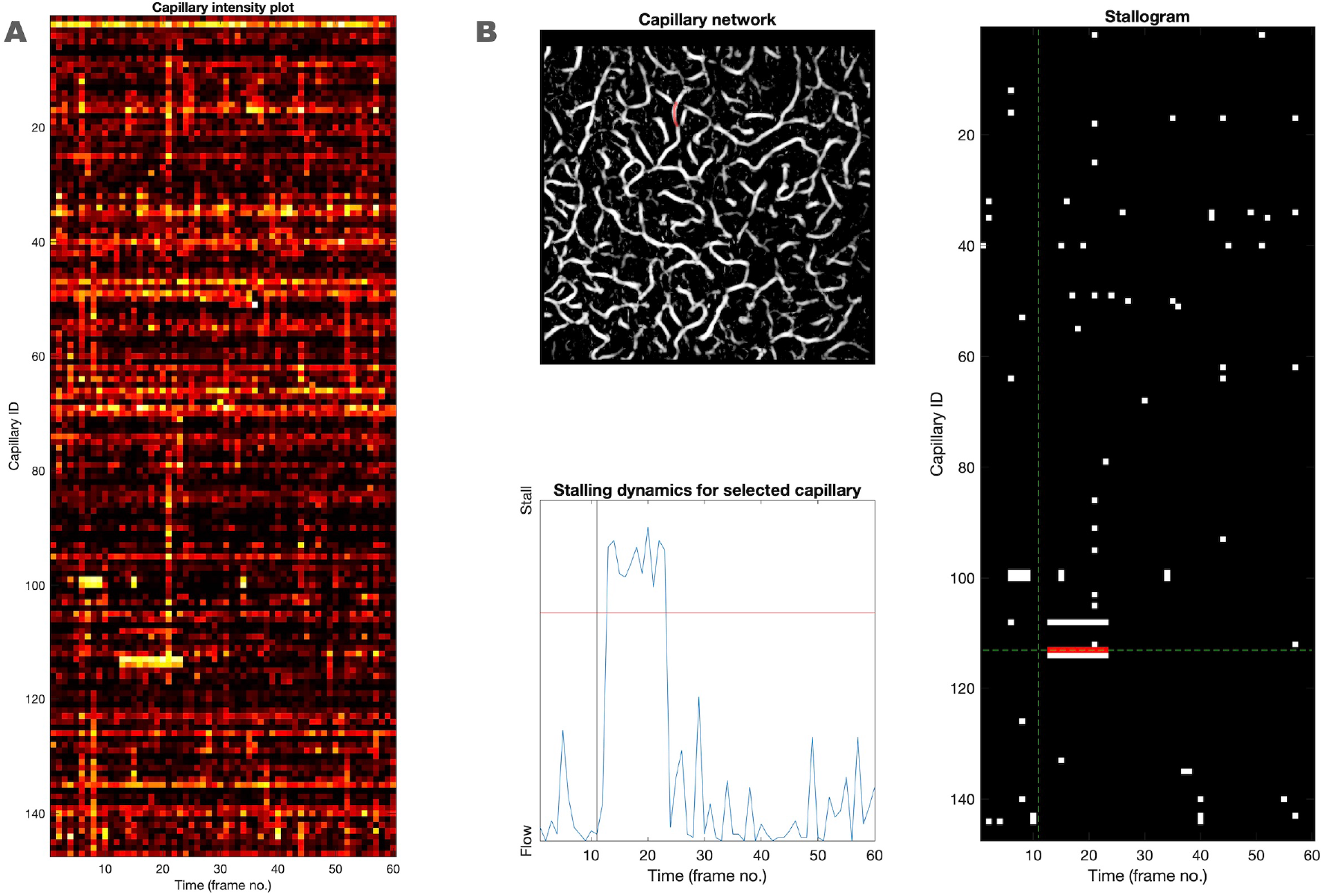
Tracking segments for identification of stalls. **A)** An un-thresholded stallogram. The colour coding reflects the image intensity at each identified capillary segment (ID). Lighter colours correspond to low values of flow, *i*.*e*., potential stalls. **B)** The GUI for adjusting the threshold of stalled segments. *Left, top*: The MIP of the selected frame (adjusted using the left and right arrow keys), showing an identified capillary in red. *Left, bottom*: Intensity profile (1 – intensity) of the capillary, where low and high intensity translate to the presence and absence of flow, respectively. The red vertical line is the threshold, which is moved up or down with the {2, w, s, x}-keys. **Right**: Thresholded stallogram showing the identified stalls through all capillaries and all frames. The horizontal and vertical line shows which capillary and frame, respectively, are shown in the images on the left. The up and down arrow keys are used to move between capillaries

**Figure 4.**
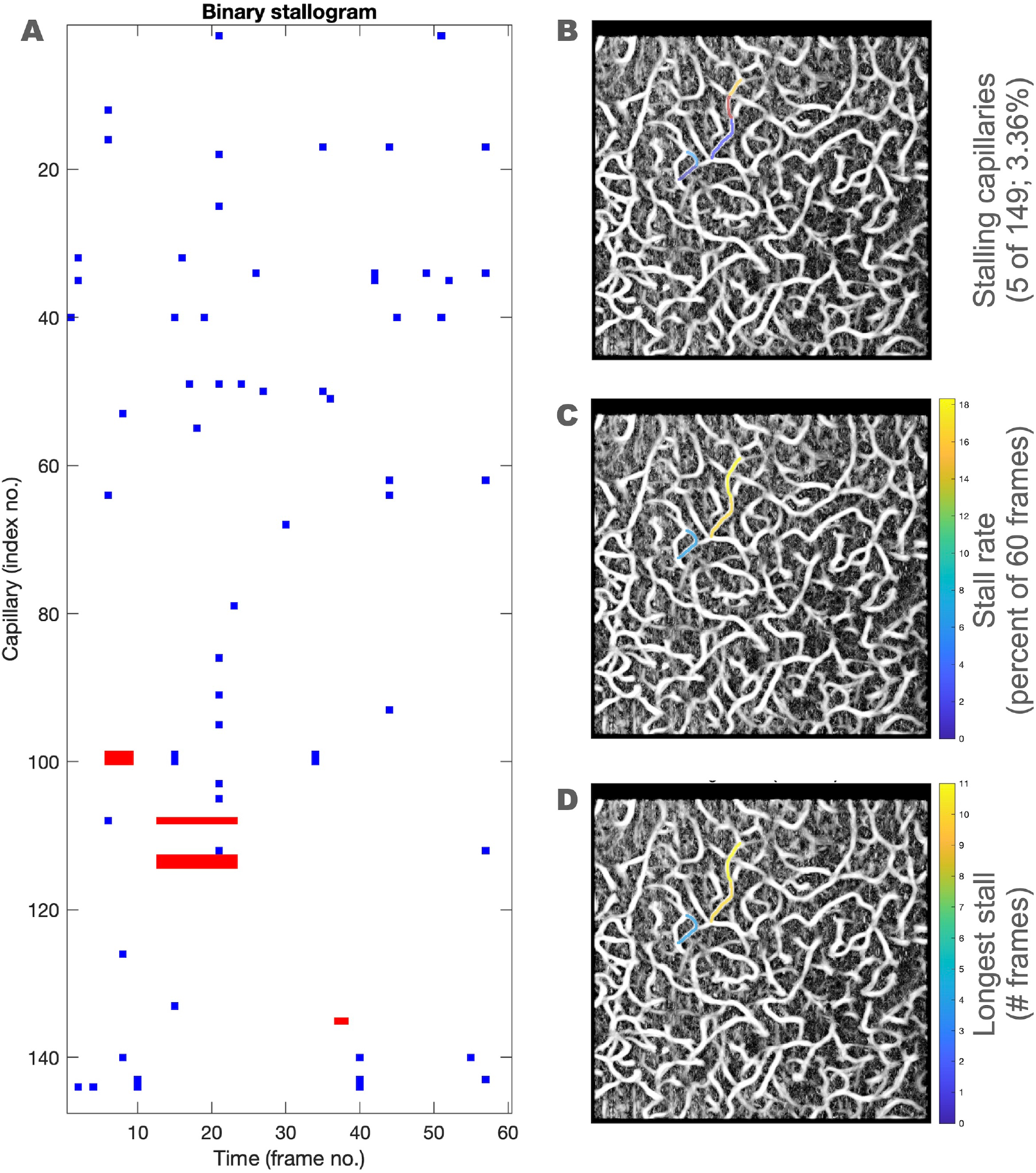
Example of results. **A)** The final stallogram after manual adjustment of stall thresholds (Figure 3B). Stalls lasting for 2 or more frames are shown in red, whereas stalls of a single frame duration are shown in blue. **B)** The capillary segments exhibiting at least one stall event during the imaging session are highlighted with a random colour. **C)** The capillary segments in A are characterised by colour depending on the relative time spent in the stalled flow-state. **D)** As in C, but colour coding for the longest stall event observed in each segment.

### Data acquisition and pre-processing

We acquired the data with a commercial spectral-domain OCT system (Telesto-II, Thorlabs, Germany), a superluminescent diode (SLD) with a central wavelength of 1310 nm and a bandwidth of 170 nm. The system has an A-line sampling frequency of 76 kHz. The estimated range in the axial direction is approximately 1.6 mm. We used a 10x objective (Mitutoyo, Kawasaki, Japan) with an estimated lateral (X) and axial (Z) resolution of 1.5 μm and 3.5 μm, respectively. The cortical microvasculature was scanned with the OCT-angiography (OCTA) technique [11]. OCTA was acquired in a 600 × 600 μm window at 400 × 400 pixels in-plane. Each image volume was acquired over ∼15 seconds, during which time movement-weighted signal is accrued for 8.5 seconds followed by a ∼6 second intertrial delay needed to save the data to disk in the Nifti-1 file format. A total of 60 such *frames* were collected in ∼15 minutes.

The purpose of pre-processing is to convert the frames (ca. 1.3 GB on disk) into 3D angiograms, *i*.*e*., volumes in which voxels containing flowing blood appear brighter than those in the background. This is the most time- and computer memory-consuming step of the analysis, which can be accelerated using parallel computing. Moreover, the structure of the data files allows for memory-efficient reading of only selected ranges of the data at a time (see below). The latter feature enables running the entire workflow on a standard dual-core laptop with only 8 GB of memory. Our implementation first converts the raw OCT spectra into reflectivity profiles. Subsequently, the two B-scans are phase-corrected, and the absolute value of their difference is calculated and returned as the angiogram [11].

### Reduction of axial (depth) dimension, image enhancement, and vessel extraction

In order to achieve a reduced memory footprint, we implemented a `Matlab` class `NiiReader` for sequential reading of the Nifti-format frames (see `doc NiiReader` for more details). A `NiiReader` object can be used to read in only a single XZ-slice (2D) from disk (Figure 1A). Furthermore, an optional `zrange`-property of the object may be used to restrict the returned data in the Z-dimension. Imaging systems that produce files in other formats may use `NiiReader` as a starting point for creating their own reader class. `NiiReader` enabled the creation of a simple graphical user interface (GUI) to identify the focus plane. Figure 1B illustrates clearly the limited effective data range. The GUI enables the definition of a restricted Z-range in which the data effectively lies. The 3D angiograms slabs are then extracted from within this range, and flattened *en face* using a maximum intensity projection (MIP): `log(max(angio))`, where the `max`-operation is applied over the Z-dimension of `angio`, and `log` is the natural logarithm.

Figure 1C-E shows the MIPs of individual frames while implementing various artefact removal procedures in sequence. OCTA images may contain acquisition artefacts which are seen as bright vertical lines in the MIP-image. To remove such stripe-artefacts, we first applied a Wavelet-based reduction technique developed in [12] (see `RemoveStripesVertical`). A dyadic Daubechies wavelet (`‘db2’`) was applied to the MIPs over 20 scales. At each scale, vertical stripes are reduced by a damping factor of 1 voxel standard deviation. Finally, the modified wavelet components were reconstructed into a de-striped angiogram image, and a 2D median filter (`medfilt2`) was applied to remove residual noise (see Figure 1C).

We employed a Frangi Vesselness filter to enhance “tubular”, vessel-like structures in the image [13]. We utilised an open-source implementation published in [14]. We applied the function `FrangiFilter2D.m` with the following parameters (manually tuned to provide robust results):

~~~
cfg_opts.FrangiScaleRange = [1, 3]; % Range of sigmas used, default [1 8]
cfg_opts.FrangiScaleRatio = 2; % Step size between sigmas, default 2
cfg_opts.FrangiBetaOne = 0.5; % Frangi correction constant, default 0.5
cfg_opts.FrangiBetaTwo = 10; % Frangi correction constant, default 15
~~~

Figure 1D illustrates the effect of the vessel-highlighting Frangi filter. Lastly, to facilitate later vessel characterisation, we enhanced contrast by applying histogram equalisation (`histeq`) onto the clean TIF frame and flooring all values less than 70% of maximum to zero (`imadjust`) (see Figure 1E).

### Identification of individual capillary segments

As a consequence of pre-processing and cleaning, each dataset is reduced to a set of 60 two-dimensional TIF images representing a snapshot of flow in the capillary bed of the imaged focus volume. Filtering artefacts at the edges of the imaging plane were mitigated by masking out a rim of 5 voxels at the XY-perimeter. An acquisition-related distortion was particularly visible in the X-dimension (see top of panel 1C), which led us to increase the masked-out segment to the first 20 voxels (see Figure 2). Both parameters should be set to reflect the properties of the dataset being analysed. Furthermore, to compensate for in-plane motion, we applied the `imregtform`-function (using the default `‘monomodal`’ optimizer and metric options) in the following manner. We identified the single frame most “similar” to the mean of all 60 images by treating each image as a 400×400-dimensional vector, and calculating the cosine of the angle between them. The identified frame was then used as the target for image registration, and all other frames were resampled to it using `imwarp`. Note that only in-plane translation parameters can be estimated in a 2D projection.

The average of all 60 frames was binarized using `imbinarize` with the ‘`adaptive’` method, which computes a local threshold at each pixel using the local mean intensity around the neighbourhood of the pixel (Figure 2A). The binary image was then passed to `bwconncomp`, which segments the image into internally-connected components (Figure 2B). We removed those components with an area less than 100 pixels; these were deemed “noise”, or simply too small structures to quantify. The cleaned binary vessel-map was then passed to `bwskel`, which erodes the binary image to its ‘skeleton’, in which only the central ‘backbone’ of each vessel branch remains (Figure 2C). We set the parameter `‘MinBranchLength’` to 10 pixels to remove the segments appearing at the endpoints of edges (not shown).

In biological terms, a *capillary segment* can be defined as a continuous stretch of vessel between two branching points. In image processing terms, the function `edgelink` [15] splits the skeletonised capillary structure into segments, which we will henceforth refer to as ‘capillary segments’ or merely ‘capillaries’. We additionally filtered out segments shorter than 16 pixels (ca. 25 μm). Using this operational definition, our method identifies on the order of 150 capillary segments in a matter of seconds (see Figure 2D and Table 1).

**Table 1.** Stall statistics in six healthy wildtype animals.

| Table 1 – Stall statistics in six healthy wildtype animals |  |  |  |  |  |
| --- | --- | --- | --- | --- | --- |
| Animal | Number of |  | Stall rate (% of 60 frames) |  |  |
|  | capillaries | stalls | mean | min | max |
| **WT01** |  |  |  |  |  |
| ROI 1 | 148 | 1 | 6.67 | 6.67 | 6.67 |
| ROI 2 | 149 | 5 | 11 | 3.33 | 26.67 |
| **WT03** |  |  |  |  |  |
| ROI 1 | 149 | 0 | 0 | 0 | 0 |
| ROI 2 | 154 | 2 | 13.33 | 3.33 | 23.33 |
| **WT04** |  |  |  |  |  |
| ROI 1 | 152 | 0 | 0 | 0 | 0 |
| ROI 2 | 168 | 3 | 8.89 | 3.33 | 18.33 |
| **WT05** |  |  |  |  |  |
| ROI 1 | 151 | 1 | 3.33 | 3.33 | 3.33 |
| ROI 2 | 159 | 1 | 6.67 | 6.67 | 6.67 |
| **WT06** |  |  |  |  |  |
| ROI 1 | 156 | 8 | 4.79 | 3.33 | 6.67 |
| ROI 2 | 156 | 5 | 6 | 3.33 | 10 |
| **WT07** |  |  |  |  |  |
| ROI 1 | 149 | 5 | 13.33 | 6.67 | 18.33 |
| ROI 2 | 149 | 5 | 11 | 3.33 | 26.67 |

### Tracking segments for identification of stalls

The objective of our analysis is to identify time points (frames) in which individual capillaries *stall, i*.*e*., when identified segments in the image *disappear*. We thus proceeded by adaptively binarizing each cleaned and motion-corrected MIP frame. The capillary segments identified in the previous step were first dilated with a 3 × 3 pixel kernel (`imdilate`), after which they were used to mask the frame data. The average value in the mask was collated for each frame, and could be visualised as a line in an `n_frames x n_capillaries` image, as shown in Figure 3A. In fact, we plot the value `1-mean(mask)`, rendering time points in which the segment is *not* present (stalled) bright, and refer to such an image as a *stallogram*.

We created a simple interactive GUI for manually adjusting the threshold level of a stall for each identified capillary segment (see Figure 3B). An initial threshold of 0.67 is used to generate an initial binary stallogram of flow (0) and stall (1) epochs. The user then adjusts the threshold for each identified capillary by determining a value corresponding to ‘true’ stalls, in contrast to noise-induced fluctuations and movement. It is also possible to mark entire frames as ‘bad’—these can be omitted when calculating stall statistics.

### Post-processing of stallograms

Information on stall events are saved as binary stallograms. From such images it is trivial to extract physiologically relevant parameters by simply ‘counting the ones’ in the rows and columns (see `runLengthEncodeRows.m`). The function `filter_stallogram.m` allows the user to filter out stalls with duration shorter than a user-defined limit prior to calculation of statistics. In the present example analysis, a ‘true’ stall was defined to have a minimum duration of 2 frames (∼30 seconds).

## Results

Figure 4A illustrates the binary stallogram obtained by applying the pipeline described in Methods. Figure 4B-D demonstrate visualisations of stall statistics, which can be readily produced for each stallogram. Figure 4B: Capillaries exhibiting at least one stall during the imaging session. Figure 4C: Stall rate, *i*.*e*., the proportion of the total imaging time a capillary is in the stalled state. Figure 4D: Longest stall duration. The statistics were calculated for stalls of at least 2 frame-duration, shown in red in Figure 4A.

Table 1 lists the stall statistics obtained by the authors. We encourage the reader interested in applying the technique on their own data to begin by reproducing our findings on the dataset available at https://osf.io/me6hf/.

## Discussion

Manual analysis workflows present several challenges to the pursuit of generalizable and reproducible research. On the one hand, the user makes explicit categorical (*e*.*g*., ‘good’ vs. ‘bad’ data segments) and ordinal (*e*.*g*., stall threshold) parameter choices that impact the summary statistics (“results”) reported. On the other hand, long sequences of complex data transformations obscure the impact of individual parameters, making it difficult to ascertain the robustness of the final product of an analysis pipeline. To address both these issues, we have here implemented a workflow in which diagnostic interactive visualisations are generated at each stage of analysis. Furthermore, we segmented our pipeline into a sequence of scripts in which the parameters relevant to each step are set in a transparent fashion.

### Methodological choices and rationale

Any random acquisition artefacts, such as the vertical stripes in our data, are relatively easy to average out if the objective is to create a static image of the vasculature. However, when detecting stalled capillaries, the time resolution is critical, and the stall information would be lost when averaging the angiograms. We therefore implemented several interventions to eliminate noise artefacts. Note, however, that the combined effect of de-striping and vessel-emphasis in frames with strong artefacts is to render such frames blank (not shown), thus leading to potential over-estimation of the number of stalls. Such commission errors can only be avoided by manually determining whether a data point must be discarded due to artefacts.

In regards to reducing the 3D volumes to 2D projections, we acknowledge that due to the complicated three-dimensional organisation of vessels in our imaging volume, and the ‘flattening’ of information content by the MIP, our method cannot achieve true parity with the biological definition of a capillary segment. In the case of two or more capillaries being on top of each other in the 3D slab, they will fuse together in the 2D projection, which may affect the result. However, by limiting the axial depth to 20 pixels, we minimize this risk while still capturing ∼150 capillaries in the 2D projection.

Our method reduces the amount of data requiring manual processing by more than three orders of magnitude from >70 GB to only 10 MB. Adequate data reduction is a critical step of implementing human-friendly analysis pipelines that present the user with only the most relevant information. Excluding the (unsupervised) maximum intensity projection-phase of analysis, our method enables users to obtain immediate feedback (within seconds) on the effects of their parameter choices. We found that with experience and judicious use of the visual guides available, a 60-frame acquisition could be manually thresholded in 10–15 minutes. We expect that simple instructions on how to operate the GUI, and on which image features correspond to ‘true’ stalls, will result in highly reproducible stall statistics.

When interpreting the results, we defined a limit of 2 consecutive frames for the identification of a “true” stall event. This was done partly to avoid characterising random noise artefacts as stall events, and partly because the data was collected from awake animals. Unanaesthetised animals, even when trained to lie still throughout the scanning session, can move. As mentioned above, only in-plane motion can be compensated for. Out-of-plane motion will confound determination of stalls from cases in which the capillary in question moves in and out of the imaging slab. Movements of these well-trained animal occur as infrequent short bursts lasting no longer than a few seconds, i.e., less than a frame (SKF, personal observation). In this example the limit was defined based on observations during acquisition. However, a motion sensor could be used to specifically eliminate frames, in which motion occurs.

### Relationship to previous work

To our knowledge, we provide here the first complete and documented analysis pipeline for MIP-based identification of capillary stall events. We note that one other preprint manuscript describes a 2D skeletonization and masking algorithm used to detect capillary stalls [16]. However, their description of methods is not sufficient to determine the extent of possible overlap. Similarly to ours, their algorithm involves a manual verification step to ensure only “true” stall events are counted in the statistics.

Stefan and Lee [17] developed deep convolutional neural networks (CNNs) for the automatic quantification of 3D OCTA images. They trained a sequence of CNNs to enhance, segment and finally graph OCTA images in a fully automated manner. The final step produces 3D vascular networks, which could potentially be used to estimate stalled flow statistics, though it was not attempted in the paper.

### Limitations

The binary nature of a stall requires setting a threshold for when features in the image correspond to an actual cessation of flow in the capillary, and when they are due to experimental noise. The `edgelink`-based identification of segments relies on 2D projections only, which will occasionally lead to the apparent fusing of two biological segments with independent flow dynamics. It will appear as if only a fraction of an automatically identified capillary is stalled, which is biologically impossible. This is one of the main reasons it is not possible to set a pixel value threshold to automatically segment a raw stallogram into a binary representation of stall events.

Another consequence of the MIP and subsequent segmentation steps is that the capillary network estimates are strongly biased towards segments that are aligned parallel to the cortical surface. Diving and surfacing vessels appear as dots in the 2D images, and are excluded from analysis by the use of minimum branch length filters. OCTA based on the intrinsic contrast mechanism of red blood cell flux was found to be severely negatively biased [18], which further validates only analysing vessels that have prominent in-plane projections.

Capillary stall events are rare in the normally functioning microcirculation of healthy animals, but may become more frequent in various disease states. Despite its limitations, we envision that the methods described here will be particularly useful in intervention studies, in which the parameter-of-interest is a manifest *change* in stalling statistics. We emphasise that our method does not purport to capture all capillaries. Rather, our aim was to obtain a large and representative sample for the calculation of various population statistics.

### Potential future developments

Although capillary segments can be defined on OCT angiograms in an automatic fashion, the determination of a threshold image intensity for stall classification remains a manual, time-consuming and potentially biased process. However, once a sufficiently large dataset has been labelled manually, it may be possible to employ supervised machine learning to train a classifier that is able to perform the task, and thus to create a fully automatic workflow.

## Author contributions

Conceived and designed the methods: SKF and CJB. Performed experiments: EGJ. Analyzed data: SKF and CJB. Wrote the paper: SKF and CJB. Contributed to the revision of the manuscript: EGJ and KRD.

## Acknowledgements

The authors wish to thank Leif Østergaard, Frédéric Lesage and Paul-James Marchand for fruitful discussions.

